# Synaptic properties of layer 6 auditory corticothalamic inputs in normal hearing and noise-induced hearing loss

**DOI:** 10.1101/2025.07.08.663770

**Authors:** Yanjun Zhao, Brandon Bizup, Thanos Tzounopoulos

**Affiliations:** Pittsburgh Hearing Research Center, Department of Otolaryngology, University of Pittsburgh, Pittsburgh, PA 15261, USA

## Abstract

Layer 6 corticothalamic neurons (CTs) provide strong feedback input that is crucial to perception and cognition in normal and pathological states; however, the synaptic properties of this input remain largely unknown, especially in pathology. Here, we examined the synaptic properties of CT axon terminals in the medial geniculate body (MGB), the auditory thalamus, in normal hearing mice and in a mouse model of noise-induced hearing loss. In normal hearing mice, we found that the synaptic strength of CTs to the core-type ventral subdivision of the auditory thalamus (MGv), which mainly conveys rapid sensory information, is stronger than the synaptic strength of CTs to the matrix-type dorsal subdivision of the auditory thalamus (MGd), which likely conveys higher-order internal state information. This is due to increased functional release sites (*n*) in CT→MGv compared to CT→MGd synapses. After noise trauma, we observed enhanced short-term facilitation in CT→MGd but not CT→MGv synapses. Our findings reveal a previously unknown mechanism of short-term synaptic plasticity after noise-induced hearing loss via which CTs enhance the throughput of matrix-type thalamus, likely to improve perceptual recovery via higher-order contextual modulation.

## Introduction

Cortico-thalamo-cortical circuits are critically involved in perception and consciousness (Guo et al., 2017; Homma et al., 2017; Voigts et al., 2020; Shepherd and Yamawaki, 2021), and thus provide a crucial network for studying synaptic mechanisms underlying perception and recovery of perception after sensory organ damage. In the cortico-thalamo-cortical loop, the reciprocity of projections between cortical and thalamic areas allows for two or more brain regions to concurrently stimulate and be stimulated by each other. This reciprocal, hierarchical and parallel processing (Ji et al., 2016) supports the synchronization and the constant bottom-up and top-down processing that allows for perception and conscious processing of sensory stimuli, such as an auditory or visual scene (Tononi and Edelman, 1998; Tononi et al., 1998; Jones, 2001; Llinas et al., 2005; Sherman, 2007; Edelman and Gally, 2013; Sherman, 2016; Shepherd and Yamawaki, 2021). Cortical projections to the thalamus are mediated by two main different classes of cortical neurons: layer (L) 6 corticothalamic (CT) neurons and L5b pyramidal tract neurons. Here, we will investigate the synaptic organization and properties between CTs and thalamic neurons and the plasticity mechanisms that alter these properties after noise-induced hearing loss (NIHL).

In the visual, somatosensory and auditory cortex, CT projections to the thalamus adjust the gain of the upper cortical layers (Olsen et al., 2012), as well as the tuning precision, synchrony, and gating of thalamic cells, as required by ongoing behavioral and coding demands (Mease et al., 2014; Guo et al., 2017; Clayton et al., 2021; Ibrahim et al., 2021). As such, CTs can shape both the overall gain and the temporal dynamics of cortical sensory responses (Mease et al., 2014). In the auditory cortex (AC), activation of CTs is crucial in gating AC population activity by desynchronizing neurons in the medial geniculate body (auditory thalamus, MGB) (Ibrahim et al., 2021) and biasing sound perception towards enhanced detection at the expense of discrimination or vice versa (Guo et al., 2017). Auditory cortex CT neurons project exclusively to the MGB, but the differences in innervation and synaptic properties between CTs and the subdivisions of the MGB are not fully understood. Moreover, how these properties change after noise-induced cochlear damage remains completely unknown.

Given the importance of CT→MGB projections in auditory perception and the prevalence of perceptual deficits noise-induced hearing loss (NIHL), it is important to understand the plasticity of CT→MGB synapses after noise trauma. Following cochlear damage, despite reduced peripheral auditory input to the brain, the neuronal activity of AC principal neurons is recovered or even enhanced due to compensatory plasticity mechanisms (Qiu et al., 2000; Auerbach et al., 2014; Chambers et al., 2016; Kumar et al., 2023). This plasticity is associated with a remarkable recovery in perceptual thresholds, but other aspects of more complex auditory processing are not compensated (Caras and Sanes, 2015; Chambers et al., 2016; Guo et al., 2017; Kumar et al., 2023). Moreover, aberrant forms of this compensatory plasticity can be associated with tinnitus and hyperacusis (Auerbach et al., 2014; Henton et al., 2023). Therefore, understanding the underlying mechanisms of plasticity following loss of peripheral input is crucial for highlighting strategies that might mitigate hearing loss and hearing loss-related disorders.

Although recent studies on AC layer (L) 2/3 revealed a cell-type specific synaptic, intrinsic and circuit plasticity among principal neurons (Kumar et al., 2023), the plasticity mechanisms in deeper layer neurons, such as the CTs, and their synapses remain largely unknown. To study the precise synaptic mechanisms in the auditory CT→MGB pathway in normal hearing and hearing loss, we used a combination of patch-clamp and anatomical techniques in a NIHL model in transgenic mice (Ntsr1-Cre). In these mice, we can selectively target and activate CT axon terminals and thus explore the potential differences in specific CT synapses on the different subdivisions of the MGB in normal and pathological hearing. Our results reveal synapse-specific short-term plasticity in the CT→MGB pathway after hearing loss, which can dynamically gate selective thalamic throughput, likely to meet the challenging perceptual demands after hearing loss.

## Materials and Methods

### Animals

Ntsr1-Cre mice (GENSAT, B6.FVB(Cg)-Tg(Ntsr1-Cre)GN220Gsat/Mmcd) were used for all experiments. All mice were handled in concordance with the National Institute of Health guidelines and approved by the Institutional Animal Care and Use Committee at the University of Pittsburgh. Experiments and analyses were blinded to the noise or sham exposure condition.

### Stereotaxic injections

Ntsr1-Cre mice (P28-35) were anesthetized with inhaled isoflurane (induction: 3 % in oxygen, maintenance: 1.5 % in oxygen) and secured in a stereotaxic frame (Kopf, Tujunga, CA). Core body temperature was maintained at ∼37 °C with a heating pad and eyes were protected with ophthalmic ointment. A small craniotomy was made for intracranial injections in the right AC (Lambda (mm); a/p: 0.0, m/l: 4.0, d/v:-1.0, w/ needle angled 30o), with a micromanipulator (Kopf).

To label and record thalamocortical neurons in the MGB, we injected ∼0.1 µl red retrograde fluorescence latex microspheres (retrobeads, Lumafluor) in the right AC. The retrobeads were pressure injected (25 psi, 10-15 msec duration) from capillary pipettes (Drummond Scientific) with a Picospritzer (Parker–Hannifin). To express ChR2 in the L6 corticothalamic pathway, we injected AAV9-EF1a-double floxed-hChR2-EYFP (titer: 2.1e^13^ vg/mL, 1:10dilution in PBS, Addgene Cat. No: 20298) in the right AC over the course of 3 minutes, using a syringe pump (World Precision Instruments, Sarasota, FL). The pipette was maintained in place for 20 minutes, then withdrawn carefully.

### Brain slice electrophysiological recordings

Patch-clamp whole-cell recordings were performed 3-4 weeks after AAV viruses and retrobead injections. Mice (P50-70) were first anesthetized with isoflurane and then immediately decapitated. Brains were rapidly removed and coronal slices (300 μm) containing the right AC and MGB were prepared in a cutting solution at 1° C using a Vibratome (VT1200 S; Leica). The cutting solution contained (in mM): 2.5 KCl, 1.25 NaH_2_PO_4_, 25 NaHCO_3_, 0.5 CaCl_2_, 7 MgCl_2_, 7 Glucose, 205 sucrose, 1.3 ascorbic acid, and 3 sodium pyruvate (pH 7.4, ∼300 mOsm, bubbled with 95% O_2_/5% CO_2_). The slices were immediately transferred and incubated at 34°C in a holding chamber for 40 min before recording. Slices were then stored at room temperature prior to recording. The incubation and recording ACSF contains the following (in mM): 125 NaCl, 2.5 KCl, 26.25 NaHCO_3_, 2 CaCl_2_, 1 MgCl_2_, 10 glucose, 1.3 ascorbic acid, and 3 sodium pyruvate (pH 7.4, ∼300 mOsm, bubbled with 95% O_2_/5% CO_2_). The recording solution was maintained at 34°C with an inline heating system (Warner TC324-B). Whole-cell recordings in voltage-and current-clamp mode were performed using a MultiClamp-700B amplifier equipped with Digidata-1440A A/D converter and Clampex (Molecular Devices). Data were sampled at 10 kHz and Bessel filtered at 4 kHz using MultiClamp 700B Commander. For voltage-clamp recordings, borosilicate pipettes (3–5 MΩ, World Precision Instruments) were filled with cesium-based internal solution containing (in mM): 126 CsCH_3_O_3_S, 4 MgCl_2_ 10 HEPES, 4 Na_2_ATP, 0.3 TrisGTP, 10 Tris-phosphocreatine, and 3 sodium ascorbate (pH ∼7.4, Osmolarity ∼300 mOsm). For current-clamp recordings, pipettes were filled with a potassium-based intracellular solution containing (in mM): 113 K-gluconate, 9 HEPES, 4.5 MgCl_2_, 4 Na-ATP, 0.3 Tris-GTP, 14 Tris phosphocreatine, 0.1 EGTA. Pipette capacitance was compensated. Series and input resistance were determined by giving a-5-mV hyperpolarizing voltage step for 50 msec in voltage-clamp mode from holding potential at-70 mV, not corrected for junction potential, and were monitored throughout the experiments. Neurons with either more than 20% of change in series or input resistance throughout the experiment or with series resistance > 25MΩ were excluded from data analysis. The images in Figures 1C-E and 2B were captured with Qcapture software and were processed with ImageJ and Adobe illustrator.

**Figure 1.**
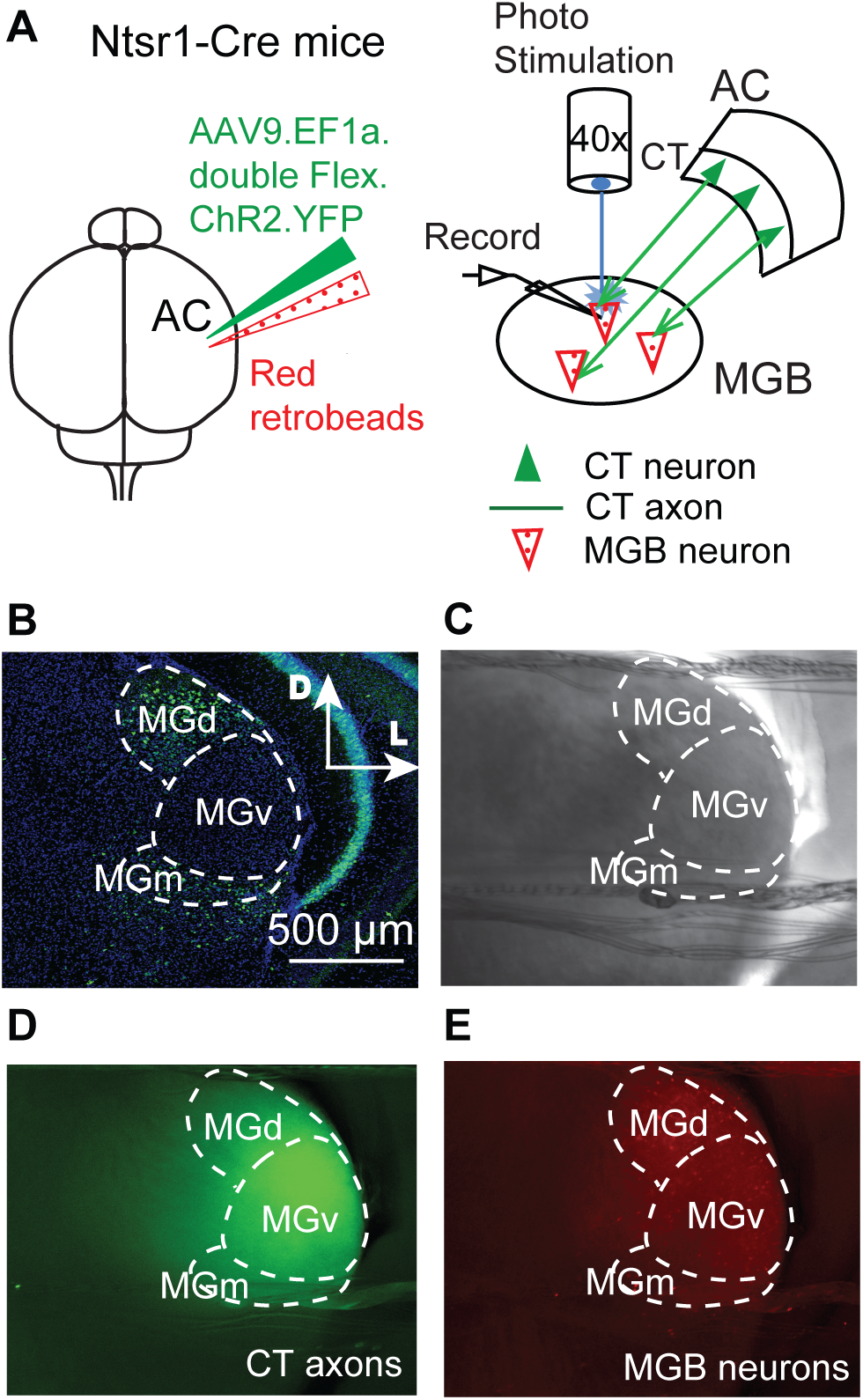
Experimental approach and MGB subdivision identification. A, Left: Schematic illustration of stereotaxic injections of retrobeads to label thalamocortical neurons (MGB neurons, red) and viral vectors (AAVs) for expression of ChR2 in corticothalamic (L6 CT) inputs in the MGB (green). All in Ntsr1-Cre mice. Right: Schematic illustration of whole-cell recordings, including CT ChR2-expressing terminals projecting to MGB principal neurons (thalamocortical neurons), which were labelled retrogradely with red retrobeads injection in AC. Photo-evoked EPSCs were recorded from MGB neurons in response to blue light stimulation. B, Representative MGB image of coronal brain slice from Ntsr1-Cre mice showing distinct MGB subdivisions based on calbindin immunostaining (green). The dorsal (MGd) and medial (MGm) MGB are strongly immunoreactive to calbindin staining. The ventral (MGv) MGB shows little immunoreactivity to calbindin staining. D: dorsal; L: lateral C, Representative image of an acute coronal brain slice that contains MGd, MGv, and MGm in bright-field (4x). Scale bar is the same as in B. D, Representative MGB image (4x) of coronal section illustrating ChR2-expressing CT projections in the MGB. Scale bar is the same as in B. E, Representative image (4x) red labelled MGB neurons, by retrobeads injected in the AC. Scale bar is the same as in B.

### Optogenetic stimulation and strontium quantal events (Sr^2+^-mEPSCs)

CT terminals that project to MGB were evoked by optogenetic stimulation, by a blue LED light source (470 nm, Thorlabs) through a 40x microscope objective lens. Recordings were targeted to the red retrobeads labeled MGB neurons. Baseline light-evoked excitatory postsynaptic currents (EPSCs) were evoked at 0.1 Hz stimulation frequency with a 10 ms light pulse duration. We initially tried to use the light intensity required to elicit a stable, maximal plateau response, but for most recordings we could not reach a plateau response even at maximal light stimulation. Therefore, for all voltage-clamp recordings, except for minimal stimulation recordings, we used maximal light stimulation. For minimal stimulation recordings, the light intensity stimulation was adjusted to produce a response failure rate > 40%. At this intensity, 100 trials were collected per cell with 0.1 Hz stimulation frequency. The average amplitude of minimal stimulation responses (minimal EPSCs) was calculated. Short-term plasticity was obtained by delivering a train of 10 stimuli of 5 ms duration each at 1 Hz, 5 Hz, and 10 Hz and with an interval of 20 s between trains. For current-clamp recordings (Figure 8), we sometimes had to avoid maximal stimulation intensity to prevent spiking during the train. Paired-pulse ratio (PPR) was calculated from the first two pulses of the 10 Hz train. Strontium quantal events were recorded using a modified Sr^2+^-ACSF solution, which contained in mM: 125 NaCl, 2.5 KCl, 26 NaHCO_3_, 4 SrCl_2_, 4 MgCl_2_, 15 glucose, 1.3 ascorbic acid, and 3 sodium pyruvate, pH 7.4, ∼300 mOsm, oxygenated w/ 95% O_2_-5% CO_2_. Slices were incubated in this solution for 30min prior to recording. All the other conditions were kept the same as described earlier. The same cutting solution, Cs^+^-based intracellular solution, and the same light activation were used. Quantal events were analyzed (Clampfit 11.2) from a 400 ms window beginning 100 ms after light stimulus (Oliet et al., 1996; Kouvaros et al., 2020). Event detection was optimized by pre-processing the traces with a lowpass Gaussian filter (1kHz). We then included events (Sr^2+^-mEPSCs) with rise time <3 ms and amplitude> 5 pA.

### Noise Induced Hearing Loss (NIHL)

Mice (P40-60) were placed in a customized 5×4 box in a sound-isolated acoustic chamber for noise or sham exposure (NE, or SE). The noise-exposed mice were exposed bilaterally to an octave band (8-16kHz) noise at 100dB SPL for 2 hours. Sham-exposed mice underwent the same exposure protocol, but without the presence of noise.

### Auditory Brainstem Responses (ABRs)

ABRs from littermate mice (P40-70) are measured from the left ear, right before SE or NE, then measured again 1d or 10d after SE or NE, immediately followed by electrophysiological recordings. Mice were placed on a heating pad (∼37°C) in a sound-attenuating chamber (ENV-022SD; Med Associates) and anesthetized under isoflurane (3% induction / 1.5% maintenance, in oxygen). ABRs were taken by placing subdermal electrodes at the vertex of the skull (active), under the left ear (reference), and under the right ear (ground). We recorded ABRs by presenting broadband clicks (1 ms duration, 0 – 80 dB SPL in 10 dB steps) and tones (0 – 80 dB SPL in 10 dB steps, 10, 12, 16, 20, 24, 32 kHz frequencies) (Kouvaros et al., 2020; Marinos et al., 2021). The sound clicks and tones were delivered through a plastic tip placed in the left ear canal at a rate of 18.56 per second with a MF1 speaker (Tucker-Davis Technologies). The speaker was calibrated with a ¼-inch microphone (4954-B, Bruel and Kjaer) using a 1 kHz, 94 dB sound calibrator standard (Type 4231, Bruel and Kjaer) as described previously (Marinos et al., 2021). For each stimulus, the evoked response was averaged from 512 trials with bandpass filtering the waveform between 300 and 3000 Hz. Data acquisition and analyses were performed using Rz6 processor and BioSigRP software. ABR threshold was defined as the lowest stimulus intensity that generated a wave I response.

### Tissue Preparation for Immunohistochemistry

Following electrophysiological recordings, brain slices were post-fixed overnight at 4°C in 4% paraformaldehyde in 1X phosphate buffered saline (PBS), pH 7.4. Slices were washed three times for 10 minutes in 1X PBS and then either entered immunohistochemistry immediately or placed in 1X PBS + 0.01% Sodium Azide at 4°C for storage.

### Immunostaining

Slices were permeabilized in 1X PBS containing 0.3% Triton-X100 (PBST) 3 times for 20 minutes each and then blocked in PBS + 5% normal goat serum (blocking buffer) at room temperature for 2 hours. Slices were then incubated in blocking buffer containing primary antibodies against calbindin (1:1000, Mouse IgG1 anti-calbindin, Swant CB300) for 72h at 4°C. Following primary antibody incubation, slices were washed 3 times in PBST for 20 minutes each. Slices were then incubated in blocking buffer containing the secondary antibody (1:1000, Goat anti-mouse IgG1 Alexa Fluor 647, Invitrogen A-21240) overnight at 4°C. Following secondary antibody incubation, slices were washed three times in PBST for 20 minutes each and three times in 1X PBS for 10 minutes each. Slices were mounted onto slides and coverslipped (Prolong Gold Antifade Mounting Medium).

### Confocal Image Acquisition and Analysis

Confocal fluorescent imaging was performed on a Leica Stellaris 5 confocal microscope using a 20X objective and using the program LASX (Leica microsystems). The approximate area of the slice containing the MGB was identified using the anti-calbindin staining. All images were acquired using the same acquisition settings. Stitched composite images were generated by capturing 2 to 4 z-stacks (0.5 mm steps, 1X zoom, pinhole: 1.0 Airy unit) of the MGB to capture the entire z-axis of the slice. Fields of view with 10% x-y overlap were captured to encompass the area of interest. The subdivisions of the MGB were determined using histological landmarks from the Allen Mouse Brain Atlas (http://mouse.brain-map.org/static/atlas) and anti-calbindin staining. The ventral portion of the MGv was distinguished from the MGd and MGm based on the anti-calbindin staining, as calbindin labels neurons that are located primarily in the dorsal and medial portions of the MGB (Lu et al., 2009; Kouvaros et al., 2023). In addition to the anti-calbindin staining, the boundaries of MGm were further estimated based on the location of the rhinal fissure, the brachium of the superior colliculus, and the shape of the hippocampal CA3 and dentate gyrus. Regions of interest (ROIs) were drawn in LASX to delineate the MGv, MGd, and MGm. To measure ChR2-eYFP fluorescence intensity, the fluorescence intensity analysis function of LASX was used in each ROI. For comparisons between SE and NE mice, average fluorescence values were normalized to the SE group. To normalize the fluorescence values, for each MGB subdivision, the average SE intensity was calculated, and each individual slice was normalized against this average value, such that the SE group average was set to 1.0.

### Statistics

Prism 9 Graphpad was used for statistical analysis. For statistical comparisons between two independent groups that passed the Shapiro-Wilk normality test, we used unpaired t-tests. The Mann-Whitney test was used for non-normally distributed data. For comparisons between multiple groups, one-way, two-way or two-way ANOVA with repeated measures were used, with post-hoc Bonferroni’s correction. Significance levels are denoted as *P < 0.05, **P < 0.01, ***P < 0.001. For detailed values and statistical tests for all figures, see Tables 1-3. Group data are presented as mean ± SEM.

**Table 1.**
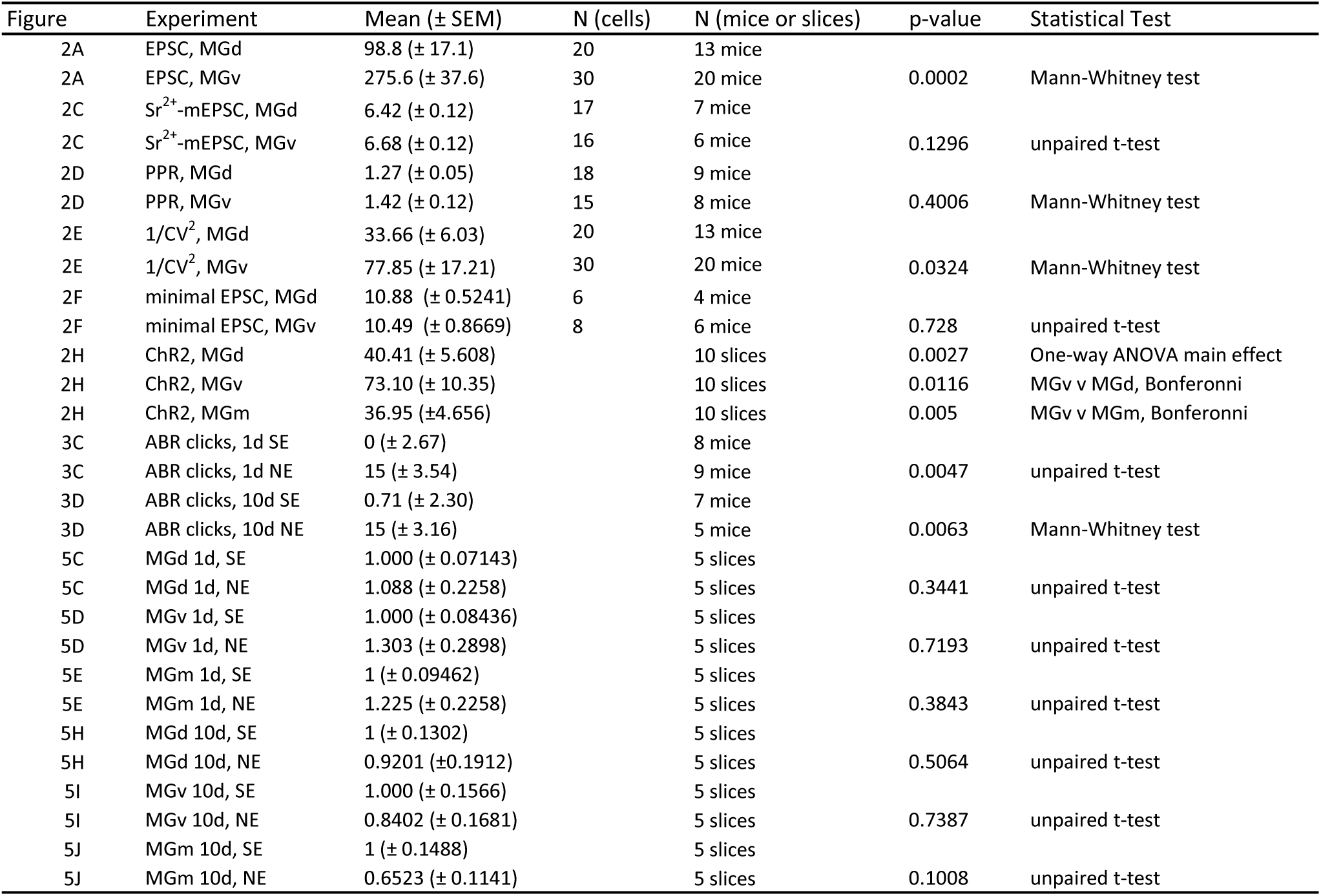
Statistical values for Figures 2,3,5.

## Results

To study the synaptic properties and organization of layer (L) 6 auditory corticothalamic (CT) inputs onto MGB neurons (CT→ MGB), we used Ntsr1-Cre transgenic mice (Figure 1A), which selectively express Cre recombinase in CTs (CT neurons, green triangle, Figure 1A). Previous studies have confirmed that all Ntsr1+ neurons in the auditory cortex (AC) are in L6 and are CTs (Guo et al., 2017). To evaluate the CT→MGB synaptic properties and organization, we selectively expressed channelrhodopsin (ChR2) in CTs (CT axons, green line, Figure 1A, D), by injecting Cre-dependent ChR2 AAV viral vector into the AC. To label MGB neurons that project to the AC (thalamocortical MGB neurons red, in Figure 1A,E), we injected red microsphere retrobeads into the AC. We then performed whole-cell recordings from acute MGB-containing brain slices (Figure 1C). We patched onto thalamocortical MGB neurons and recorded light-evoked EPSCs in response to photo-stimulating CT terminals that expressed ChR2. To evaluate the localization of the MGB neurons in the different MGB subdivisions, we used an anti-calbindin antibody, which preferentially labels the medial (MGm) and dorsal (MGd) but not the ventral (MGv) MGB (Lu et al., 2009; Kouvaros et al., 2023) (Figure 1B). Due to the smaller area, neuronal visibility and borderline localization of the MGm in our acute brain slices, we were unable to obtain enough whole-cell recordings from that area. We therefore focused our electrophysiological studies on MGv and MGd neurons.

### CTs form functional synapses that are stronger in CT→MGv compared to CT→MGd synapses, due to higher *n* in CT→MGv

We found that the average EPSC peak amplitude was significantly smaller in the CT→MGd synapses compared CT→MGv synapses (Figure 2A). Whereas the observed changes in synaptic strength based on optogenetics approaches are subject to potential variable infection efficiency per mouse/brain area, in Figure 2B we show a representative example of recordings from different subdivisions within the same slice, further supporting the effect we observed by averaging EPSCs from MGB neurons from different mice and brain slices.

**Figure 2.**
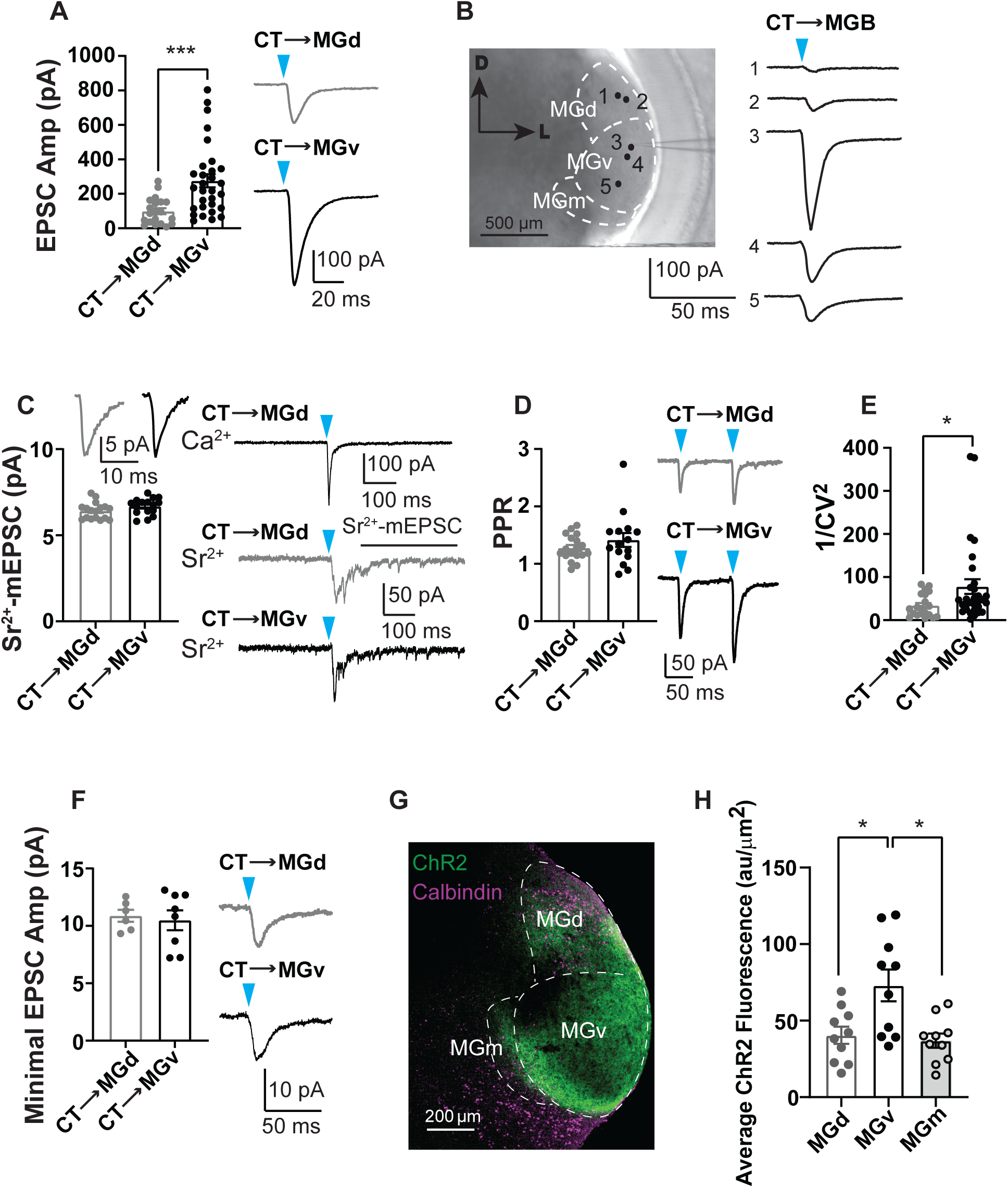
CT→MGv synapses are stronger than to CT→MGd synapses, due to higher *n* in CT→MGv synapses. A, Average light-evoked EPSC amplitude in CT→MGd and CT→MGv synapses (left) and representative traces (right). P = 0.0002 (Mann-Whitney test, CT→MGd: 20 cells; CT→MGv: 30 cells). Data for CT→MGd and CT→MGv are pooled from the 1d SE and 10d SE of CT→MGd and CT→MGv (Figure 4 A,F) recordings, respectively. B, CT→MGB EPSC traces from the same slice. Recorded neurons are numbered from 1 to 5, in order from MGd to MGv. C, Average light-evoked quantal EPSCs (Sr^2+^-mEPSCs) in CT→MGd and CT→MGv synapses (left bottom), and representative average traces (left top). Right: The top right trace (black) is a control trace from Ca^2+^-containing ACSF; the middle and bottom traces are from Sr^2+^-containing ACSF. The arrowhead indicates the onset of light stimulus. The solid line indicates the 400 ms time window beginning 100ms after light stimulus that was used to analyze the amplitude of Sr^2+^-mEPSC. P = 0.1296 (unpaired t test, CT→MGd: 17 cells; CT→MGv: 16 cells). Data for CT→MGd and CT→MGv are pooled from the 1d SE and 10d SE of CT→MGd and CT→MGv (Figure. 4C,H) recordings, respectively. D. Average paired pulse ratio (PPR) in CT→MGd and CT→MGv synapses (left) and representative traces (right). P = 0.4006 (Mann-Whitney test, CT→MGd: 18 cells; CT→MGv: 15 cells). Data for CT→MGd and CT→MGv are pooled from the 1d SE and 10d SE of CT→MGd and CT→MGv recordings (Figure. 4D,I), respectively E, Average 1/CV^2^ in CT→MGd and CT→MGv synapses. P = 0.0324 (Mann-Whitney test, CT→MGd: 20 cells; CT→MGv: 30 cells). Data for CT→MGd and CT→MGv are pooled from the 1d SE and 10d SE of CT→MGd and CT→MGv recordings (Figure. 4E,J), respectively. F, Average minimal EPSC amplitude (left) and representative averaged minimal EPSC traces from one cell (right) from CT→MGd (grey) and CT→MGv (black) synapses. P = 0.728 (unpaired t test, CT→MGd: 6 cells; CT→MGv: 8 cells). G, Representative image of coronal brain slice showing calbindin immunostaining (magenta) and ChR2 fluorescence (green). H, Quantification of ChR2 immunofluorescence intensity in the different MGB subdivisions. P = 0.0027 (One-way ANOVA, MGd: 10 slices; MGv: 10 slices; MGm: 10 slices). Data for MGd and MGv are pooled from the 1d SE and 10d SE of MGd and MGv (Figure. 5C,D,H,I) respectively. Detailed statistical values are listed in Table 1.

To investigate the mechanisms underlying the difference in synaptic strength between CT→MGd and CT→MGv synapses, we employed various assays to evaluate postsynaptic and/or presynaptic mechanisms. First, we collected and analyzed the evoked quantal events in Sr^2+^ (Sr^2+^-mEPSCs) (Oliet et al., 1996; Kouvaros et al., 2020). Replacing Ca^2+^ with Sr^2+^ in the ACSF (Methods) desynchronizes the evoked neurotransmitter release, thus allowing the analysis of quantal events from the stimulated synapses (Oliet et al., 1996). We found that the average amplitude of the quantal events (*q*) was not different between CT→MGv and CT→MGd synapses (Figure 2C), supporting that the observed differences in EPSC size are not associated with changes in *q*.

Because we did not detect any changes in *q*, we next investigated whether presynaptic mechanisms might account for the differential EPSC amplitude between CT→MGd and CT→MGv synapses. Presynaptic mechanisms were evaluated by analyzing the paired pulse ratio (PPR) and 1/CV^2^ (Faber and Korn, 1991; Zucker and Regehr, 2002). An increase in PPR would indicate a decrease in the probability of neurotransmitter release (*p*), whereas a decrease in PPR would indicate an increase in *p*. An increase in 1/CV^2^ would indicate an increase in the number of functional releasing sites (*n*) and/or an increase in *p*, whereas a decrease in 1/CV^2^ would indicate a decrease in *n* and/or *p*. We found that both synapses showed facilitation and PPR was not different between CT→MGd and CT→MGv synapses (Figure 2D), indicating that *p* is not different in these synapses. We found a significantly greater 1/CV^2^ in CT→MGv compared to CT→MGd synapses (Figure 2E), likely indicating a higher *n* in L6CT→MGv. Because *n* is the product of individual synapse strength and number of synapses activated during the stimulus, we used minimal stimulation to determine the synaptic strength after activation of a single axon. We found that minimal response was not different between CT→MGv compared to CT→MGd synapses, suggesting that synaptic strength in response to a single axon stimulation is not different between CT→MGv and CT→MGd synapses (Figure 2F). To further evaluate these findings, we quantified the average ChR2 immunofluorescence intensity in the MGd, MGv and MGm nuclei. We found that the average immunofluorescence intensity was significantly higher in MGv than MGd and MGm (Figure 2G,H), suggesting a higher density of CT→MGv axons than CT→MGd axons. Together, these results support that although either *q* or the strength of individual cortical inputs are not different between CT→MGv and CT→MGd synapses, a higher density of axons in CT→MGv synapses underlies the increased synaptic strength of CT→MGv compared to CT→MGd synapses.

### No changes in baseline synaptic transmission in either CT→MGv or CT→MGd synapses after noise-induced hearing loss (NIHL)

Next, we assessed whether noise trauma affects synaptic properties in either CT→MGv and/or CT→MGd synapses. To answer this question, we used an NIHL mouse model, where we exposed mice bilaterally to an octave band (8-16 kHz) noise at 100 dB SPL for 2 h (Figure 3A, Methods, (Kumar et al., 2023)). As control, we used sham-exposed (SE) mice, which were treated identically to the noise-exposed (NE) mice but without noise presentation (Methods). To evaluate hearing thresholds, we measured auditory brainstem responses (ABRs), which represent synchronized neural activity in response to sound stimuli from the auditory nerve to the inferior colliculus along the auditory brainstem. The first wave originates from the type I auditory nerve fibers and reflects hearing thresholds (Buchwald and Huang, 1975; Kiang et al., 1976). ABRs were collected before, 1day (d) after, and 10d after NE or SE (Figure 3B). ABR threshold shifts were calculated by subtracting the pre-exposure ABR threshold from the post-exposure ABR threshold. A positive ABR threshold shift indicates hearing loss, as evidenced by an increase in hearing thresholds after noise exposure. We found that ABR thresholds were elevated at 1d and remained elevated at 10d after NE (Figure 3C-E). In contrast, ABR thresholds remained unchanged in the 1d or 10d in SE mice (Figure 3C-E), supporting that our mouse model captures hearing loss that lasts at least for 10d.

**Figure 3.**
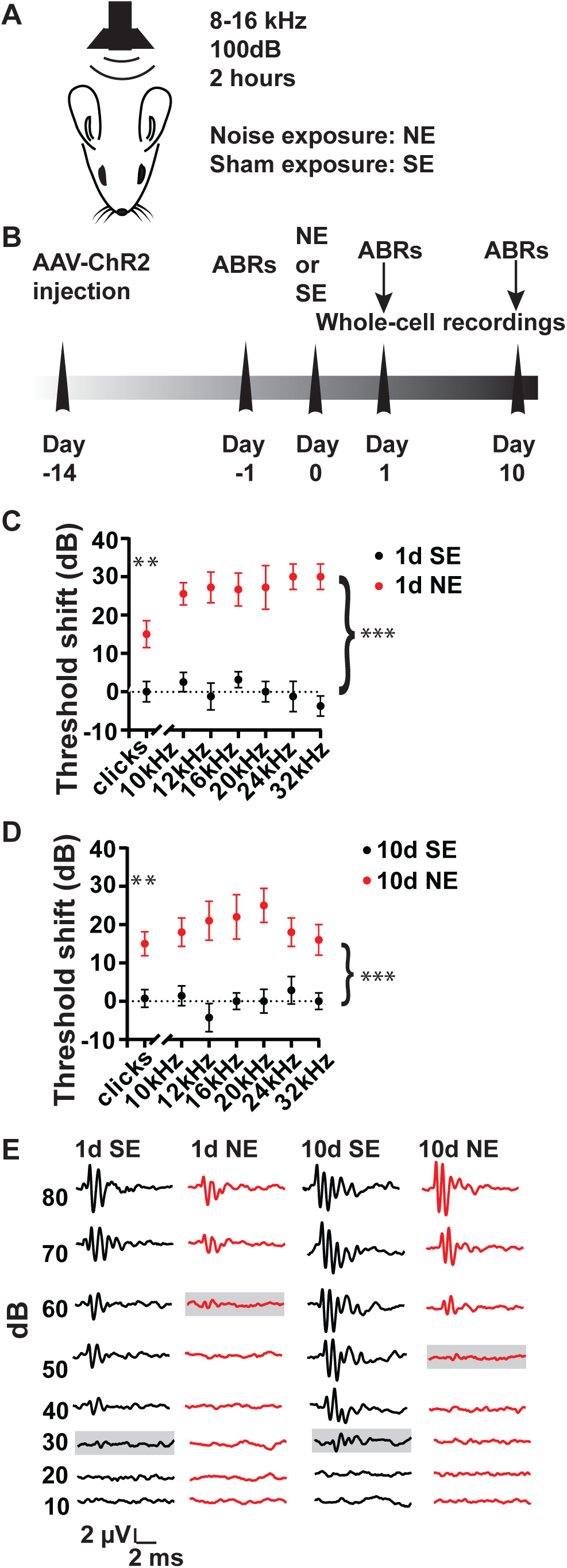
Mouse model of noise-induced hearing loss (NIHL) A, Noise exposure paradigm. Mice were bilaterally exposed to either an 8-16kHz sound at 100 dB for 2 hours or sham exposure. B, Timeline of experimental design. C, Average ABR threshold shifts (clicks and pure tones) from 1d SE and 1d NE mice. Clicks, P = 0.0047 (unpaired t test, 1d SE: 8 mice; 1d NE: 9 mice); pure tones, p < 0.0001 (effect of exposure, 2-way ANOVA, 1d SE: 8 mice; 1d NE: 9 mice). D, Average ABR threshold shift (clicks and pure tones) from 10d SE and 10d NE mice. Clicks, P = 0.0063 (Mann-Whitney test, 10d SE: 7 mice; 10d NE: 5 mice); pure tones, p < 0.0001 (effect of exposure, 2-way ANOVA, 10d SE: 7 mice; 10d NE: 5 mice). E, Representative ABR traces in response to clicks, from 1d SE, 1d NE, 10d SE, and 10d NE mice. Highlighted traces indicate the ABR thresholds. Detailed statistical values are listed in Tables 1,2.

We used this NIHL mouse model to examine the potential changes in the properties of CT→MGv and CT→MGd synapses. Whole-cell recordings of MGB neurons were performed from acutely prepared brain slices 1d or 10d after NE or SE. To study the synaptic properties of the CT-MGB pathway after hearing loss, we patched onto MGB neurons located at either MGd or MGv and recorded light-evoked EPSCs by photo-stimulating CT terminals that expressed ChR2, as described previously. We found no changes in the EPSC amplitude 1d or 10d after NE in either CT→MGd or CT→MGv synapses (Figure 4A,F). We did not observe any changes in membrane input resistance (Figure 4B,G), quantal size (Figure 4C,H), 1/CV^2^ (Figure 4D,I) between NE vs. SE in either CT→MGd or CT→MGv synapses (Figure 4). However, we found an increase in PPR 1d after NE in CT→MGd synapses (Figure 4E), suggesting a decrease in *p*, which recovers by 10d after NE (Figure 4J).

**Figure 4.**
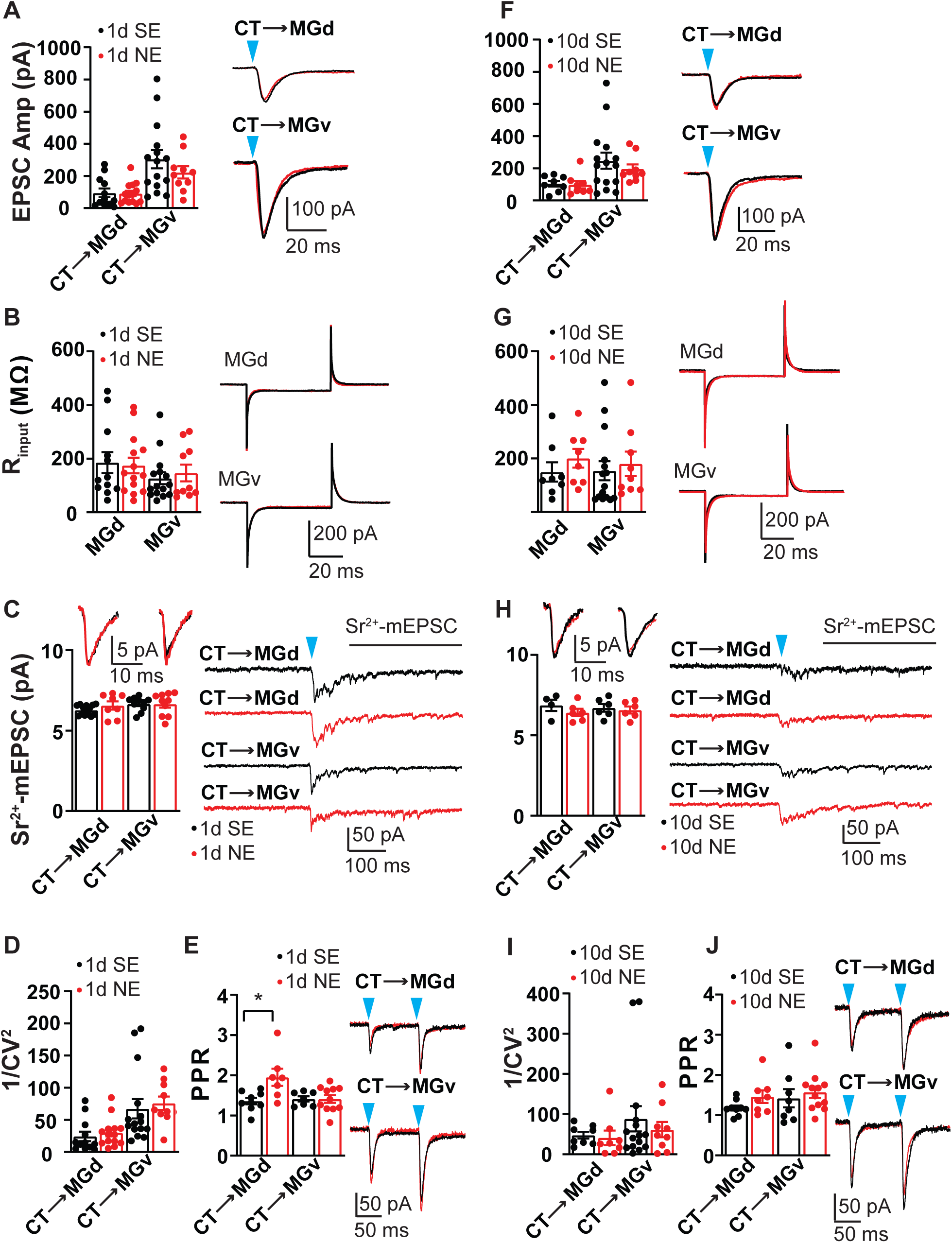
No changes in baseline synaptic transmission in either CT→MGv or CT→MGd synapses after NIHL. A, Average ESPC peak amplitude in 1d SE and NE in CT→MGd and CT→MGv synapses (left), and representative traces (right), P = 0.2898 (effect of exposure, 2-way ANOVA, MGd 1d SE: 12 cells; MGd 1d NE: 14 cells; MGv 1d SE: 15 cells; MGv 1d NE: 10 cells). B, Average membrane input resistance in 1d SE and 1d NE in MGd or MGv neurons (left), and representative traces (right). P = 0.8898 (effect of exposure, 2-way ANOVA, MGd 1d SE: 12 cells; MGd 1d NE: 14 cells; MGv 1d SE: 15 cells; MGv 1d NE: 10 cells). C, Average Sr^2+^-mEPSC amplitude in 1d SE vs 1d NE in CT→MGd and CT→MGv synapses (bottom left) and representative average traces (top left). The arrowhead indicates the onset of light stimulus. The solid line indicates the 400 ms time window beginning 100ms after light stimulus that was used to analyze the amplitude of Sr^2+^-mEPSCs. P = 0.4407 (effect of exposure, 2-way ANOVA, MGd 1d SE: 13 cells; MGd 1d NE: 8 cells; MGv 1d SE: 10 cells; MGv 1d NE: 10 cells). D, Average 1/CV^2^ in 1d SE vs 1d NE in CT→MGd and CT→MGv synapses. P = 0.5186 (effect of exposure, 2-way ANOVA, MGd 1d SE: 12 cells; MGd 1d NE: 14 cells; MGv 1d SE: 15 cells; MGv 1d NE: 10 cells). E, Average PPR, in 1d SE vs 1d NE in CT→MGd and CT→MGv synapses (left), and representative traces (right). P = 0.0193 (effect of exposure, 2-way ANOVA, MGd 1d SE: 9 cells; MGd 1d NE: 7 cells; MGv 1d SE: 7 cells; MGv 1d NE: 11 cells). F, Average ESPC peak amplitude in 10d SE and 10d NE in CT→MGd and CT→MGv synapses (left), and representative traces (right). P = 0.5133 (effect of exposure, 2-way ANOVA, MGd 10d SE: 8 cells; MGd 10d NE: 8 cells; MGv 10d SE: 15 cells; MGv 10d NE: 9 cells). G, Average membrane input resistance in 10d SE and 10d NE in CT→MGd and CT→MGv synapses (left), and representative traces (right). P = 0.3498 (effect of exposure, 2-way ANOVA, MGd 10d SE: 8 cells; MGd 10d NE: 8 cells; MGv 10d SE: 15 cells; MGv 10d NE: 9 cells). H, Average Sr^2+^-mEPSC amplitude in 10d SE and 10d NE in CT→MGd and CT→MGv synapses (bottom left) and representative average traces (top left). The arrowhead indicates the onset of the light stimulus. The solid line indicates the 400 ms time window beginning 100ms after the light stimulus that was used to analyze the amplitude of Sr^2+^-mEPSCs. P = 0.2674 (effect of exposure, 2-way ANOVA, MGd 10d SE: 4 cells; MGd 10d NE: 6 cells; MGv 10d SE: 6 cells; MGv 10d NE: 6 cells). I, Average 1/CV^2^ in 10d SE and 10d NE in CT→MGd and CT→MGv synapses. P = 0.5502 (effect of exposure, 2-way ANOVA, MGd 10d SE: 8 cells; MGd 10d NE: 8 cells; MGv 10d SE: 15 cells; MGv 10d NE: 9 cells). J, Average PPR in 10d SE and 10d NE in CT→MGd and CT→MGv synapses (left), and representative traces (right). P = 0.1727 (effect of exposure, 2-way ANOVA, MGd 10d SE: 9 cells; MGd 10d NE: 8 cells; MGv 10d SE: 8 cells; MGv 10d NE: 12 cells). Detailed statistical values are listed in table 3.

Finally, when we quantified the ChR2 immunofluorescence intensity in MGv and MGd, we did not observe any differences between NE and SE (Figure 5), consistent with our electrophysiological results. Together, these results support that baseline synaptic transmission at low frequency (0.1 Hz) of CT activation is not altered in either CT→MGv or CT→MGd synapses after NIHL. However, we observed a change in the PPR of CT→MGd synapses when two pulses were delivered at a 10 Hz frequency (100 ms interstimulus interval). This prompted us to investigate whether there are any differences in short-term plasticity (STP) in response to different stimulation frequencies before and after noise trauma.

**Figure 5.**
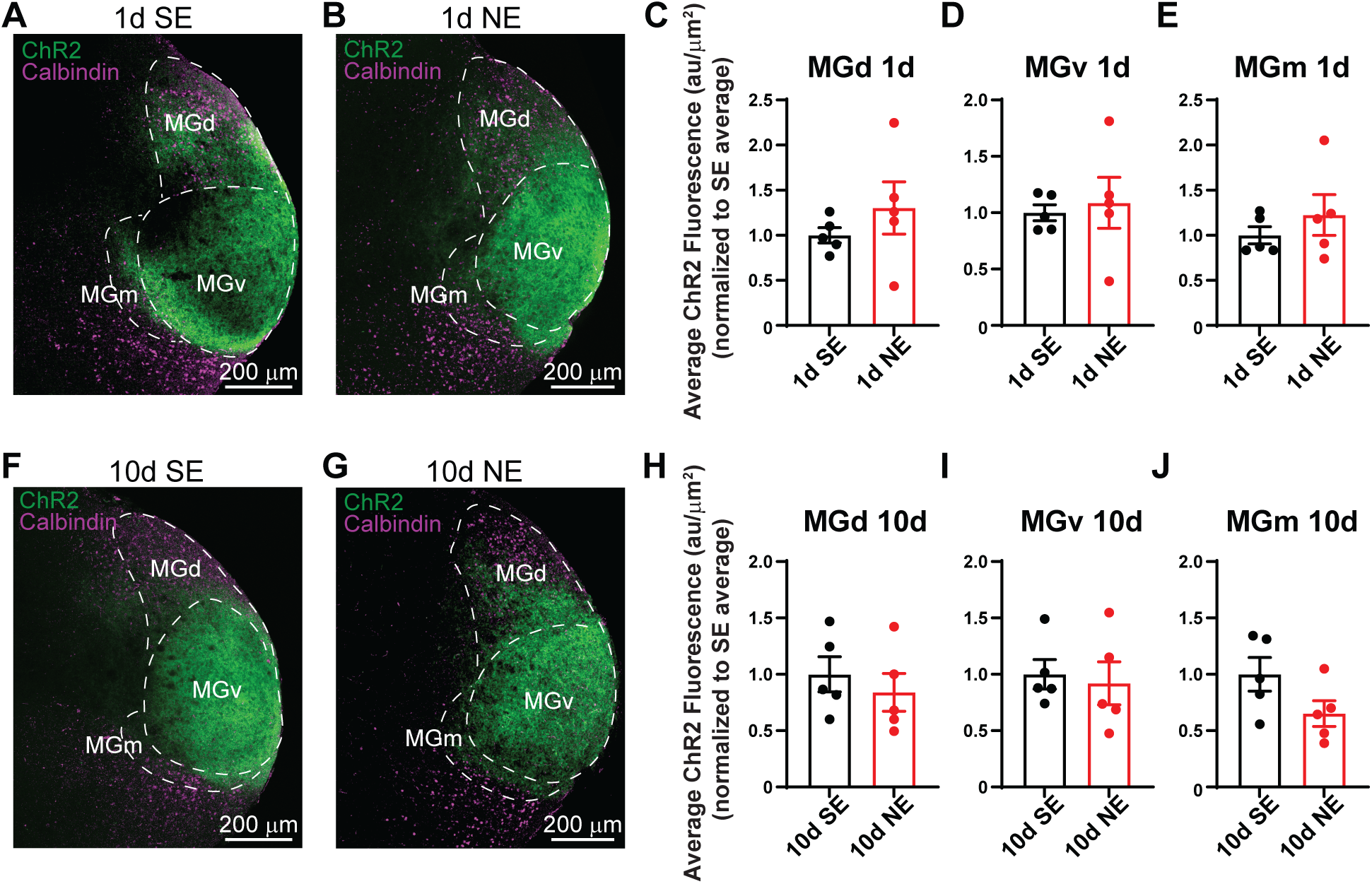
Unchanged ChR2 fluorescence intensity of CT terminals in the different MGB subdivisions after NIHL. A-B, Representative images of coronal brain slices showing calbindin immunostaining (magenta) and ChR2 fluorescence (green) from 1d SE (A) and 1 NE (B). C-E, Average ChR2 immunofluorescence intensity in the different MGB subdivisions in 1d SE and 1d NE (C, MGd; D, MGv; E: MGm). C, P = 0.3441 (unpaired t test, 1d SE: 5 slices; 1d NE: 5 slices); D, P = 0.7193 (unpaired t test, 1d SE: 5 slices; 1d NE: 5 slices); E, P = 0.3843 (unpaired t test, 1d SE: 5 slices; 1d NE: 5 slices). F-G, Representative images of coronal brain slices showing calbindin immunostaining (magenta) and ChR2 fluorescence (green) from 10d SE (F) and 10d NE (G). H-J, Average ChR2 immunofluorescence intensity in the different MGB subdivisions at 10d SE and 10d NE (H, MGd; I, MGv; J: MGm). H, P = 0.5064 (unpaired t test, 10d SE: 5 slices; 10d NE: 5 slices); I, P = 0.7387 (unpaired t test, 10d SE: 5 slices; 10d NE: 5 slices); J, P = 0.1008 (unpaired t test, 10d SE: 5 slices; 10d NE: 5 slices). Detailed statistical values are listed in table 1.

### Enhanced activity-dependent facilitation in CT→MGd, but not CT→MGv synapses after NIHL

In the somatosensory cortex, under normal (sparse, or baseline or low) frequency of CT firing, CT→ventral posterior medial nucleus (VPm, core-type thalamus) synaptic activity leads to a transient increase in thalamic neuronal firing rates, which is followed by longer period of robust suppression (Crandall et al., 2015). However, when CTs fire at moderately higher frequencies, such as 5-10Hz, thalamic neurons show enhanced spiking activity. This transformation is due to facilitation of excitation in CT → VPm synapses and a depression of thalamic reticular nucleus (TRN)→VPm inhibition (Crandall et al., 2015). This corticothalamic switch is important for sensory processing, as it acts as an activity-dependent gating mechanism of sensory input inflow to cortex. Thus, we investigated whether there is an activity-dependent facilitation in AC CT → MGd/v synapses following different frequencies of CT stimulation and whether this facilitation is affected after NIHL. To test this, we delivered a train of 10 brief blue light pulses at frequencies of 1Hz, 5Hz, and 10Hz, respectively. The peak amplitude of the evoked EPSCs was normalized to the 1^st^ peak in the train for comparison. Consistent with previous studies in somatosensory cortex (Crandall et al., 2015), we observed facilitation of both CT→MGd and CT→MGv synapses at 5 and 10 Hz, but not at 1Hz (Figures 6 and 7, SE). Moreover, we did not observe any difference in short-term dynamics between CT →MGd and CT→MGv synapses in SE. Importantly, although we did not find any changes in short-term dynamics in L6 CT→MGv synapses either 1d or 10d after NE vs. SE (Figure 7), we found increased facilitation in CT→MGd synapses in NE vs. SE both in 1d (Figure 6C) and 10d after NE (Figure 6E). To further validate the physiological significance of these findings in CT→MGd synapses, we employed current-clamp mode experiments to explore the interaction of synaptic and intrinsic properties in shaping STP. Consistent with our recordings in voltage-clamp mode, we found increased facilitation in CT→MGd synapses in NE vs. SE 1 d after exposure (Figure 8), further supporting the physiological significance of our findings. Together, these results support that peripheral hearing loss induces synapse-specific changes on STP in CT→MGB synapses, with enhanced STP in CT→MGd but not CT→MGv synapses. Importantly, these results suggest that after hearing loss, while the corticothalamic auditory CT input to the core-type (lemniscal) MGv pathway remains unchanged, there is an activity-dependent preferential enhancement of CT input to the matrix-type (non-lemniscal) MGd, supporting a compensatory mechanism that likely facilitates auditory perception after hearing loss via activation of matrix-type thalamic neurons and higher-level cortical activity (Discussion).

**Figure 6.**
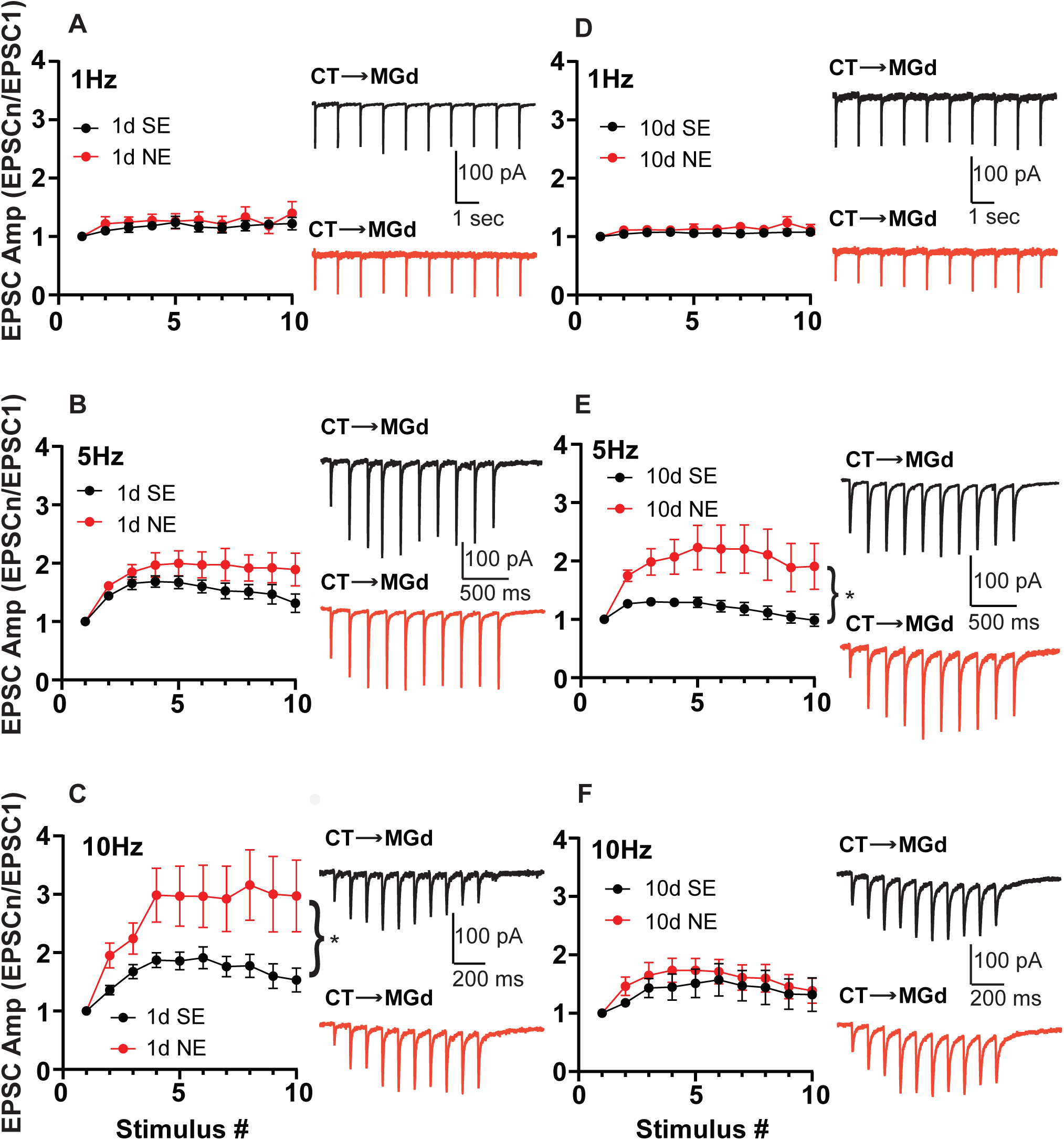
Enhanced CT→MGd activity-dependent facilitation after NIHL. A-B, Average peak amplitudes of EPSCs in the train, normalized to the peak amplitude of the first EPSC in CT→MGd synapses, in response to a train of 10 light pulses at 1 Hz (A), 5 Hz (B), and 10 Hz (C), in 1d SE (black) and 1d NE (red). A, P = 0.5266 (effect of exposure, 1d SE: 10 cells; 1d NE: 9 cells); B, P = 0.1613 (effect of exposure, 1d SE: 9 cells; 1d NE: 11 cells); C, P = 0.0265 (effect of exposure, 1d SE: 9 cells; 1d NE: 7 cells). D-F, Average peak amplitudes of EPSCs in the train, normalized to the peak amplitude of the first EPSC in CT→MGd synapses, in response to a train of 10 light pulses at 1 Hz (A), 5 Hz (B), and 10 Hz (C), in 10d SE (black) and 10d NE (red). D, P = 0.3406 (effect of exposure, 10d SE: 8 cells; 10d NE: 7 cells); E, P = 0.0239 (effect of exposure, 10d SE: 7 cells; 10d NE: 6 cells); F, P = 0.5526 (effect of exposure, 10d SE: 9 cells; 10d NE: 8 cells). Detailed statistics values are listed in table 2.

**Figure 7.**
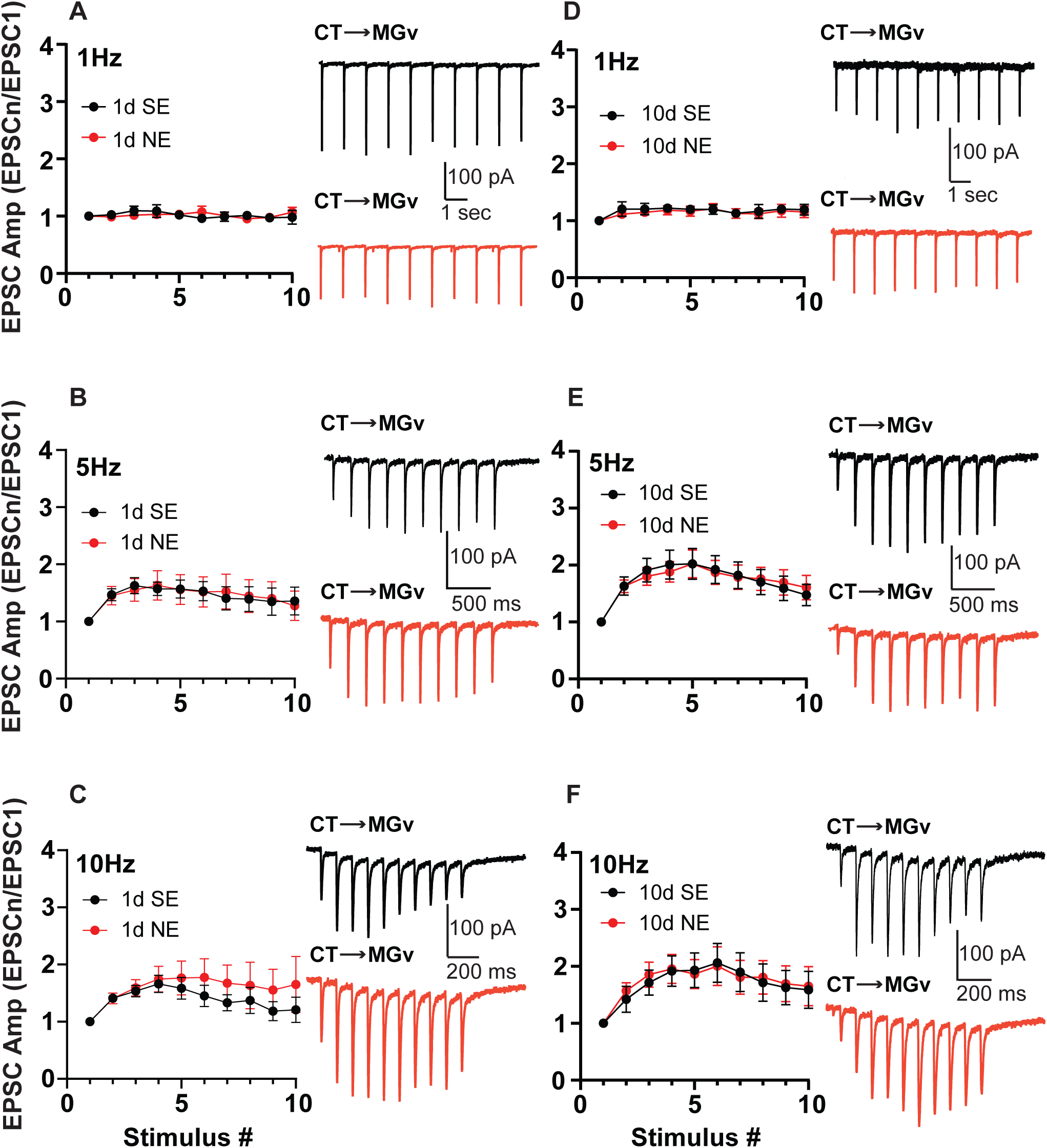
No changes in a CT→MGv activity-dependent facilitation after NIHL. A-C, Average peak amplitudes of EPSCs in the train, normalized to the peak amplitude of the first EPSC in CT→MGv synapses, in response to a train of 10 light pulses at 1 Hz (A), 5 Hz (B), and 10 Hz (C), in 1d SE (black) and 1d NE (red). A, P = 0.9964 (effect of exposure, 1d SE: 4 cells; 1d NE: 8 cells); B, P = 0.9781 (effect of exposure, 1d SE: 6 cells; 1d NE: 10 cells); C, P = 0.5555 (effect of exposure, 1d SE: 7 cells; 1d NE: 11 cells). D-F, Average peak amplitudes of EPSCs in the train, normalized to the peak amplitude of the first EPSC in CT→MGv synapses, in response to a train of 10 light pulses at 1 Hz (A), 5 Hz (B), and 10 Hz (C), in 10d SE (black) and 10d NE (red). D, P = 0.7594 (effect of exposure, 10d SE: 8 cells; 10d NE: 12 cells); E, P = 0.9921 (effect of exposure, 10d SE: 6 cells; 10d NE: 12 cells); F, P = 0.9258 (effect of exposure, 10d SE: 8 cells; 10d NE: 12 cells). Detailed statistical values are listed in table 2.

**Figure 8.**
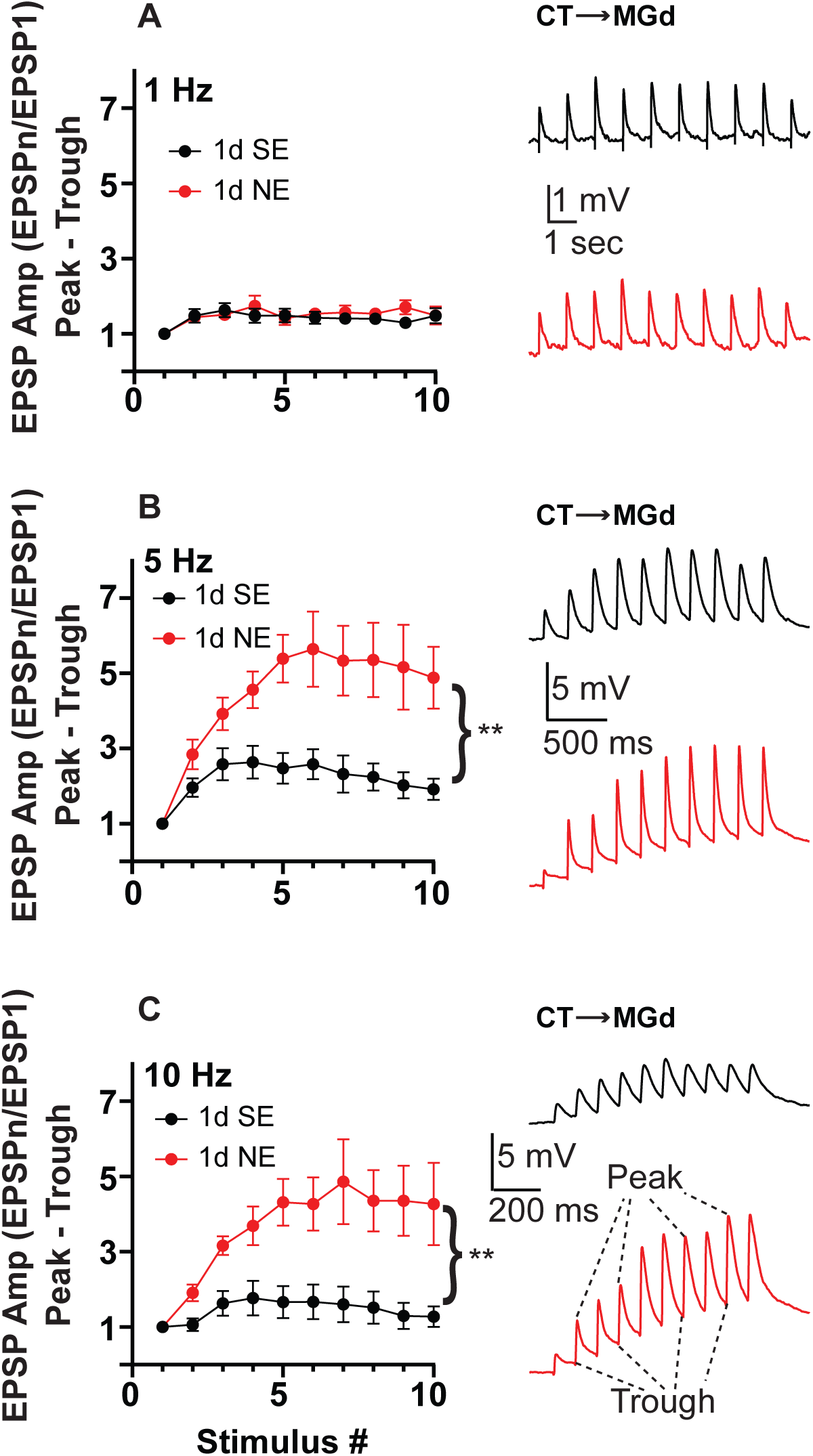
Enhanced CT→MGd activity-dependent facilitation after NIHL in current-clamp mode. A-C, Average trough-subtracted peak EPSP amplitudes, normalized to the peak amplitude of the first EPSP in CT→MGd synapses, in response to a train of 10 light pulses at 1 Hz (A), 5 Hz (B), and 10 Hz (C), in 1d SE (black) and 1d NE (red). Peak-trough value was obtained by subtracting the peak amplitude of the EPSP from the through amplitude of the preceding EPSP. The first value is not a peak-trough value, it is the normalized value of the first EPSP peak amplitude. A, P = 0.6477 (effect of exposure, 1d SE: 6 cells; 1d NE: 5 cells); B, P = 0.0102 (effect of exposure, 1d SE: 6 cells; 1d NE: 5 cells); C, P = 0.0076 (effect of exposure, 1d SE: 6 cells; 1d NE: 5 cells).

**Table 2.** Statistical values for Figures 3, 6, 7, 8.

| Figure | Experiment | Statistical test | Effect of exposure | Effect of stimulus | Interaction effect | Multiple comparison | N (cells/mice) |
| --- | --- | --- | --- | --- | --- | --- | --- |
| 3C | ABR pure tone, 1d SE vs 1d NE | 2-way ANOVA, Bonferroni | p < 0.0001 | p = 0.9948 | p = 0.6447 | 10 kHz, p = 0.0001<br>12 kHz, p < 0.0001<br>16 kHz, p < 0.0001<br>20 kHz, p < 0.0001<br>24 kHz, p < 0.0001<br>32 kHz, p < 0.0001 | 1d SE = na/8<br>1d NE = na/9 |
| 3D | ABR pure tone, 10d SE vs 10d NE | 2-way ANOVA, Bonferroni | p < 0.0001 | p = 0.8293 | p = 0.5473 | 10 kHz, p = 0.0130<br>12 kHz, p < 0.0001<br>16 kHz, p = 0.0004<br>20 kHz, p < 0.0001<br>24 kHz, p = 0.0289<br>32 kHz, p = 0.0180 | 10d SE = na/7<br>10d NE = na/5 |
| 6A | 1Hz MGd 1d SE vs 1d NE | 2-way repeated measures ANOVA; Bonferroni | p = 0.5266 | p = 0.0571 | p = 0.8175 | ns all stimuli | 1d SE = 10/5<br>1d NE = 9/6 |
| 6B | 5Hz MGd 1d SE vs 1d NE | 2-way repeated measures ANOVA; Bonferroni | p = 0.1613 | p = 0.0010 | p = 0.1208 | ns all stimuli | 1d SE = 9 /4<br>1d NE = 11/8 |
| 6C | 10Hz MGd 1d SE vs 1d NE | 2-way repeated measures ANOVA; Bonferroni | p = 0.0265 | p = 0.0013 | p = 0.0008 | ns all stimuli | 1d SE = 9 /4<br>1d NE = 7/6 |
| 6D | 1Hz MGd 10d SE vs 10d NE | 2-way repeated measures ANOVA; Bonferroni | p = 0.3406 | p = 0.1028 | p = 0.4997 | ns all stimuli | 10d SE = 8 /5<br>10d NE = 7/5 |
| 6E | 5Hz MGd 10d SE vs 10d NE | 2-way repeated measures ANOVA; Bonferroni | p = 0.0239 | p = 0.0124 | p < 0.0001 | Stimulus #1 p = 0.0258<br>ns all other stimuli | 10d SE = 7 /4<br>10d NE = 6/5 |
| 6F | 10Hz MGd 10d SE vs 10d NE | 2-way repeated measures ANOVA; Bonferroni | p = 0.5526 | p = 0.0188 | p = 0.9552 | ns all stimuli | 10d SE = 9/5<br>10d NE = 8/6 |
| 7A | 1Hz MGv 1d SE vs 1d NE | 2-way repeated measures ANOVA; Bonferroni | p = 0.9964 | p = 0.4677 | p = 0.5538 | ns all stimuli | 1d SE = 4/4<br>1d NE = 8/5 |
| 7B | 5Hz MGv 1d SE vs 1d NE | 2-way repeated measures ANOVA; Bonferroni | p = 0.9781 | p = 0.0334 | p = 0.9966 | ns all stimuli | 1d SE = 6/4<br>1d NE = 10/6 |
| 7C | 10Hz MGv 1d SE vs 1d NE | 2-way repeated measures ANOVA; Bonferroni | p = 0.5555 | p = 0.0889 | p = 0.8480 | ns all stimuli | 1d SE = 7/4<br>1d NE = 11/6 |
| 7D | 1Hz MGv 10d SE vs 10d NE | 2-way repeated measures ANOVA; Bonferroni | p = 0.7594 | p = 0.0708 | p = 0.9944 | ns all stimuli | 10d SE = 8/4<br>10d NE = 12/6 |
| 7E | 5Hz MGv 10d SE vs 10d NE | 2-way repeated measures ANOVA; Bonferroni | p = 0.9921 | p = 0.0003 | p = 0.9332 | ns all stimuli | 10d SE = 6/3<br>10d NE = 12/5 |
| 7F | 10Hz MGv 10d SE vs 10d NE | 2-way repeated measures ANOVA; Bonferroni | p = 0.9258 | p = 0.0041 | p = 0.9916 | ns all stimuli | 10d SE = 8 /4<br>10d NE = 12/6 |
| 8A | 1Hz MGd 1d SE vs 1d NE | 2-way repeated measures ANOVA; Bonferroni | p = 0.6477 | p = 0.0439 | p = 0.4431 | ns all stimuli | 1d SE = 6/5<br>1d NE = 5/3 |
| 8B | 5Hz MGd 1d SE vs 1d NE | 2-way repeated measures ANOVA; Bonferroni | p = 0.0102 | p = 0.0008 | p < 0.0001 | ns all stimuli | 1d SE = 6/5<br>1d NE = 5/3 |
| 8C | 10Hz MGd 1d SE vs 1d NE | 2-way repeated measures ANOVA; Bonferroni | p = 0.0076 | p = 0.0067 | p < 0.0001 | Stimulus #2 p = 0.0479<br>ns all other stimuli | 1d SE = 6/5<br>1d NE = 5/3 |

**Table 3.** Statistical values for Figure 4.

| Figure | Statistical test | Effect of exposure | Effect of location | Interaction effect | N (cells/mice) | Multiple comparison |
| --- | --- | --- | --- | --- | --- | --- |
| 4A, EPSC | 2-way ANOVA<br>Bonferroni | p = 0.2898 | p = 0.0001 | p = 0.3380 | MGd 1d SE = 12/6 | MGd 1d SE vs MGd 1d NE, p = 0.9999 |
|  |  |  |  |  | MGd 1d NE = 14/10 | MGd 1d SE vs MGv 1d SE, p = 0.0023 |
|  |  |  |  |  | MGv 1d SE = 15/9 | MGv 1d SE vs MGv 1d NE, p = 0.4961 |
|  |  |  |  |  | MGv 1d NE = 10/7 | MGd 1d NE vs MGv 1d NE, p = 0.1304 |
| 4B, R<sub>in</sub> | 2-way ANOVA<br>Bonferroni | p = 0.8898 | p = 0.1651 | p = 0.6199 | MGd 1d SE = 12/6 | MGd 1d SE vs MGd 1d NE, p > 0.9999 |
|  |  |  |  |  | MGd 1d NE = 14/10 | MGd 1d SE vs MGv 1d SE, p > 0.9999 |
|  |  |  |  |  | MGv 1d SE = 15/9 | MGv 1d SE vs MGv 1d NE, p > 0.9999 |
|  |  |  |  |  | MGv 1d NE = 10/7 | MGd 1d NE vs MGv 1d NE, p > 0.9999 |
| 4C, Sr<sup>2+</sup>-<br>mEPSC | 2-way ANOVA<br>Bonferroni | p = 0.4407 | p = 0.1814 | p = 0.3938 | MGd 1d SE = 13/5 | MGd 1d SE vs MGd 1d NE, p = 0.6562 |
|  |  |  |  |  | MGd 1d NE = 8/4 | MGd 1d SE vs MGv 1d SE, p = 0.3501 |
|  |  |  |  |  | MGv 1d SE = 10/4 | MGv 1d SE vs MGv 1d NE, p > 0.9999 |
|  |  |  |  |  | MGv 1d NE = 10/3 | MGd 1d NE vs MGv 1d NE, p = 0.9870 |
| 4D, 1/CV<sup>2</sup> | 2-way ANOVA<br>Bonferroni | p = 0.5186 | p = 0.0002 | p = 0.8974 | MGd 1d SE = 12/6 | MGd 1d SE vs MGd 1d NE, p > 0.9999 |
|  |  |  |  |  | MGd 1d NE = 14/10 | MGd 1d SE vs MGv 1d SE, p = 0.0367 |
|  |  |  |  |  | MGv 1d SE = 15/9 | MGv 1d SE vs MGv 1d NE, p > 0.9999 |
|  |  |  |  |  | MGv 1d NE = 10/7 | MGd 1d NE vs MGv 1d NE, p = 0.0374 |
| 4E, PPR | 2-way ANOVA<br>Bonferroni | p = 0.0193 | p = 0.0507 | p = 0.0187 | MGd 1d SE = 9/4 | MGd 1d SE vs MGd 1d NE, p = 0.0105 |
|  |  |  |  |  | MGd 1d NE = 7/6 | MGd 1d SE vs MGv 1d SE, p > 0.9999 |
|  |  |  |  |  | MGv 1d SE = 7/4 | MGv 1d SE vs MGv 1d NE, p > 0.9999 |
|  |  |  |  |  | MGv 1d NE = 11/6 | MGd 1d NE vs MGv 1d NE, p = 0.0164 |
| 4F, EPSC | 2-way ANOVA<br>Bonferroni | p = 0.5133 | p = 0.0089 | p = 0.6122 | MGd 10d SE = 8/7 | MGd 10d SE vs MGd 10d NE, p > 0.9999 |
|  |  |  |  |  | MGd 10d NE = 8/6 | MGd 10d SE vs MGv 10d SE, p = 0.1165 |
|  |  |  |  |  | MGv 10d SE = 15/10 | MGv 10d SE vs MGv 10d NE, p > 0.9999 |
|  |  |  |  |  | MGv 10d NE = 9/5 | MGd 10d NE vs MGv 10d NE, p = 0.8291 |
| 4G, R<sub>in</sub> | 2-way ANOVA<br>Bonferroni | p = 0.3498 | p = 0.8421 | p = 0.7575 | MGd 10d SE = 8/7 | MGd 10d SE vs MGd 10d NE, p > 0.9999 |
|  |  |  |  |  | MGd 10d NE = 8/6 | MGd 10d SE vs MGv 10d SE, p > 0.9999 |
|  |  |  |  |  | MGv 10d SE = 15/10 | MGv 10d SE vs MGv 10d NE, p > 0.9999 |
|  |  |  |  |  | MGv 10d NE = 9/5 | MGd 10d NE vs MGv 10d NE, p > 0.9999 |
| 4H, Sr<sup>2+</sup>-<br>mEPSC | 2-way ANOVA<br>Bonferroni | p = 0.2674 | p = 0.9917 | p = 0.5361 | MGd 10d SE = 4/2 | MGd 10d SE vs MGd 10d NE, p = 0.6401 |
|  |  |  |  |  | MGd 10d NE = 6/2 | MGd 10d SE vs MGv 10d SE, p = 0.9725 |
|  |  |  |  |  | MGv 10d SE = 6/2 | MGv 10d SE vs MGv 10d NE, p = 0.9799 |
|  |  |  |  |  | MGv 10d NE = 6/2 | MGd 10d NE vs MGv 10d NE, p = 0.9657 |
| 4I, 1/CV<sup>2</sup> | 2-way ANOVA<br>Bonferroni | p = 0.5502 | p = 0.2759 | p = 0.7108 | MGd 10d SE = 8/7 | MGd 10d SE vs MGd 10d NE, p > 0.9999 |
|  |  |  |  |  | MGd 10d NE = 8/6 | MGd 10d SE vs MGv 10d SE, p > 0.9999 |
|  |  |  |  |  | MGv 10d SE = 15/10 | MGv 10d SE vs MGv 10d NE, p > 0.9999 |
|  |  |  |  |  | MGv 10d NE = 9/5 | MGd 10d NE vs MGv 10d NE, p > 0.9999 |
| 4J, PPR | 2-way ANOVA<br>Bonferroni | p = 0.1727 | p = 0.2644 | p = 0.6737 | MGd 10d SE = 9/5 | MGd 10d SE vs MGd 10d NE, p > 0.9999 |
|  |  |  |  |  | MGd 10d NE = 8/6 | MGd 10d SE vs MGv 10d SE, p > 0.9999 |
|  |  |  |  |  | MGv 10d SE = 8/4 | MGv 10d SE vs MGv 10d NE, p > 0.9999 |
|  |  |  |  |  | MGv 10d NE = 12/6 | MGd 10d NE vs MGv 10d NE, p > 0.9999 |

## Discussion

Here, we used Ntsr1-Cre mice to study the synaptic properties of L6 CT neurons. Previous results indicate that Ntsr1+ CT neurons reside in upper L6 (L6a) and lower L6 (L6b) (Olsen et al., 2012; Chevee et al., 2018), with the L6a CTs projecting mainly to core-type and L6b CTs projecting mainly to matrix-type thalamus (Bourassa and Deschenes, 1995; Llano and Sherman, 2008; Chevee et al., 2018; Hoerder-Suabedissen et al., 2018; Zolnik et al., 2020; Antunes and Malmierca, 2021). Although our studies did not determine the exact sublayer origin of the CT neurons, our results showing that CTs project to MGv and MGd are consistent with previous studies from somatosensory and motor cortices, which show that Ntsr1+ CTs project to core-and matrix-type thalamus (Yamawaki and Shepherd, 2015; Chevee et al., 2018; Frandolig et al., 2019; Guo et al., 2020; Antunes and Malmierca, 2021; Shepherd and Yamawaki, 2021).

One limitation of our study is that our experiments were conducted in 2mM external Ca^2+^. While we understand that 1.2 mM external Ca^2+^ is more closely related to *in vivo* conditions (Borst, 2010; Forsberg et al., 2019), the main finding of our study showing enhanced facilitation at 5 and 10Hz CT activation in CT→MGd after noise trauma is likely underestimated in our recording conditions. An additional limitation of our study is that we have not investigated yet the mechanisms underlying the rate-dependent increase in CT →MGd synapses after noise trauma. One potential mechanism would be that noise trauma lowers the threshold for recruiting a reserve pool of vesicles (Denker and Rizzoli, 2010) during the train stimuli. Moreover, changes in the recruitment of neuromodulatory systems during trains of activity after noise trauma, such as endocannabinoid signaling (Kalappa and Tzounopoulos, 2017; Whitt et al., 2022), might be involved in the observed changes in synaptic dynamics. In futures studies, we plan to explore the precise synaptic mechanisms underlying this noise-induced plasticity in the synaptic dynamics of CT→MGd synapses.

In the somatosensory cortex, CTs project more strongly to the core-vs. matrix-type thalamocortical neurons (Guo et al., 2020), but in motor cortex CTs project stronger to matrix-vs. the core-type thalamocortical neurons (Harris and Shepherd, 2015; Shepherd and Yamawaki, 2021). However, the auditory cortical CT innervation and synaptic strength in the different subdivisions of the MGB have not been fully determined. Here, we found that the synaptic strength of CT→MGv is stronger than the synaptic strength of CT→MGd, likely due to increased *n* in CT→MGv synapses, which is consistent with the somatosensory cortex findings. One limitation of our approach is that we do not know the precise localization of CTs within the different subfields of the AC. Based on previous anatomical studies in the AC, the CT→MGv input is likely originating from A1 (Llano and Sherman, 2008), which is tonotopically organized and has relatively simple receptive fields (lemniscal pathway). The CT→MGd input is likely arising from the secondary AC (AII) and a dorsoposterior region (DP) (Llano and Sherman, 2008), both of which have weaker tonotopy and more complex receptive fields (non-lemniscal pathway).

In the somatosensory system, the effect of CT neuronal feedback on thalamus is activity-dependent. Namely, low-frequency (0.1Hz) CT stimulation leads to a small transient excitation in the somatosensory thalamus (VPm) that is followed by a strong inhibition and reduced spiking (Crandall et al., 2015). This happens because the relatively weak CT excitatory input to VPm is followed by the strong disynaptic inhibitory input via the TRN, which sends inhibition to the VPm. On the contrary, higher-frequency (5-10 Hz) CT stimulation leads to strong excitation and enhanced VPm spiking (Crandall et al., 2015). This activity-dependent switch is due to the differences in the short-term plasticity at higher frequencies of CT synapses to TRN and the primary sensory thalamic nucleus. Namely, TRN→VPm synapses exhibit short-term depression, while CT→VPm synapses exhibit short-term facilitation, thereby altering the balance of excitatory and inhibitory inputs to the VPm driven by CT neurons. The visual system displays similar dynamics (Jurgens et al., 2012; Whitt et al., 2022). Here, we found that AC also follows similar dynamics, at least in terms of the short-term plasticity of the CT→MGv and CT→MGd synapses, which show no plasticity at 1Hz (Figures 6A,D; 7A,D; 8A) but short-term potentiation at 5 and 10Hz (Figures 6B,C,E,F; 7B,C,E,F; 8B,C). Although, we did not test the short-term dynamic of TRN→MGB, our results support that sensory cortices can gate their own sensory input via an activity-dependent mechanism that controls L6 corticothalamic input.

In the sensory cortex, it has been proposed that the CT projection to core-type thalamus participates in reciprocal cortico-thalamo-cortical circuit and modulates sensory processing, while the CT projection to the matrix-type thalamus likely participates in higher-order functions, such as perception, cognition and attention (Llano and Sherman, 2008; Homma et al., 2017; Hoerder-Suabedissen et al., 2018; Zolnik et al., 2020; Antunes and Malmierca, 2021; Ibrahim et al., 2021). However, the synaptic mechanisms of these two pathways are largely unknown in the auditory system and especially in hearing loss. In this context, the most important finding of our study is that after noise-induced hearing loss we observed an enhanced facilitation at 5 and 10Hz CT activation in CT→MGd but not CT→MGv synapses. It has been known that the reduction of auditory nerve input to the brain after noise trauma induces robust plasticity in the AC, and other brain regions of the auditory system, that compensates for the reduced sensory input (Qiu et al., 2000; Auerbach et al., 2014; Chambers et al., 2016; Kumar et al., 2023). However, our findings uncover a previously unknown corticothalamic plasticity mechanism after noise trauma that highlights enhanced CT input to MGd after noise trauma. Given that MGd is a matrix-type thalamic nucleus that projects to higher order cortical areas that can generate transthalamic activation from lower to higher order cortical areas(Sherman, 2016), this mechanism might enhance perceptual recovery after noise trauma via the enhanced involvement of the MGd feedback and higher cortical areas that modulate perception and cognition (Sherman, 2016; Shepherd and Yamawaki, 2021).

Alternatively, or additionally, the noise-trauma-induced enhancement of the MGd output that terminates in the L1 apical tuft dendrites of L5 pyramidal neurons would also enhance perceptual recovery after hearing loss, according to the back-propagation activated coupling model (Jones, 1998, 2001; Llinas et al., 2005; Larkum, 2013). This model does not depend on transthalamic activation across different areas but, instead, on the integration of information from thalamocortical and corticothalamic pathways at the cellular level, namely at L5 pyramidal neurons. According to this model, enhancement of the perceptual recovery would be achieved by enhanced integration of matrix-type MGd and higher-cortex input, which conveys internal representation information onto the L1 apical dendrites of L5 pyramidal neurons (Larkum, 2013), with the core-type MGv input, which conveys external sensory information onto the basal dendrites of the same L5 pyramidal neurons (Larkum, 2013). The enhanced MGd feedback after noise trauma would likely enhance perceptual recovery via enhanced contextual modulation of perception after NIHL and thus enhancement of prediction (Jones, 1998, 2001; Super et al., 2001; Llinas et al., 2005; Larkum, 2013). Importantly, recent data support that L5 dendritic activity in the AC mostly correlates with behavioral actions rather than sensory processing (Ford et al., 2024). Thus, the noise-trauma-induced enhancement of the MGd output, which terminates in the L1 apical tuft dendrites of L5 pyramidal neurons, could also enhance perceptual recovery via enhanced behavioral rather than sensory processing performance.

It is important to note that we cannot exclude the possibility that the plasticity described here might contribute to tinnitus, where internally generated percepts, which are likely stored in our brain for predictive purposes, can be released involuntarily (Henton and Tzounopoulos, 2021). In this context, the enhanced feedback inputs after hearing loss could contribute to tinnitus. Taken together, further understanding of the synaptic mechanisms underlying plasticity in cortico-thalamo-cortical loops after peripheral trauma will not only elucidate cortical plasticity mechanisms after sensory organ damage but also holds the potential to highlight novel targets that may either enhance perceptual recovery after sensory organ damage or mitigate plasticity-related sensory processing disorders, such as tinnitus, hyperacusis and phantom limb pain.

